# Comprehensive profiling of antibody responses to the human anellome using programmable phage display

**DOI:** 10.1101/2022.03.28.486145

**Authors:** Thiagarajan Venkataraman, Harish Swaminathan, Cesar A. Arze, Sarah M. Jacobo, Agamoni Bhattacharyya, Tyler David, Dhananjay M. Nawandar, Simon Delagrave, Vinidhra Mani, Nathan L. Yozwiak, H. Benjamin Larman

## Abstract

Viruses belonging to the diverse *Anelloviridae* family represent a major constituent of the commensal human virome. Aside from their widespread prevalence and persistence in humans and their absence of detectable pathologic associations, little is known about the immunobiology of the human anellome. In this study, we employed the Phage ImmunoPrecipitation Sequencing (PhlP-Seq) assay for comprehensive analyses of antibody binding to 56 amino acid long anellovirus peptides. We designed and constructed a large and diverse “AnelloScan” T7 phage library comprising more than 32,000 non-redundant peptides representing the ORF1, ORF2, ORF3 and TTV-derived apoptosis-inducing protein (TAIP) sequences of more than 800 human anelloviruses (spanning three genera). We used this library to profile the antibody reactivities of serum samples from 156 subjects. The vast majority of anellovirus peptides were not reactive in any of the subjects tested (n=~28,000; ~85% of the library). Antibody reactive peptides were largely restricted to the C-terminal region of the putative capsid protein, ORF1. To characterize antibody responses to newly acquired anellovirus infections, we screened a longitudinal cohort of matched blood-transfusion donors and recipients. Most transmitted anelloviruses did not elicit detectable antibody reactivity in the recipient (29 out of a total of 40 transmitted anelloviruses) and the remainder demonstrated delayed reactivity (~100-150 days after transfusion). This study represents the first large-scale epitope-level serological survey of the antibody response to the human anellome.

**Graphical Abstract:** 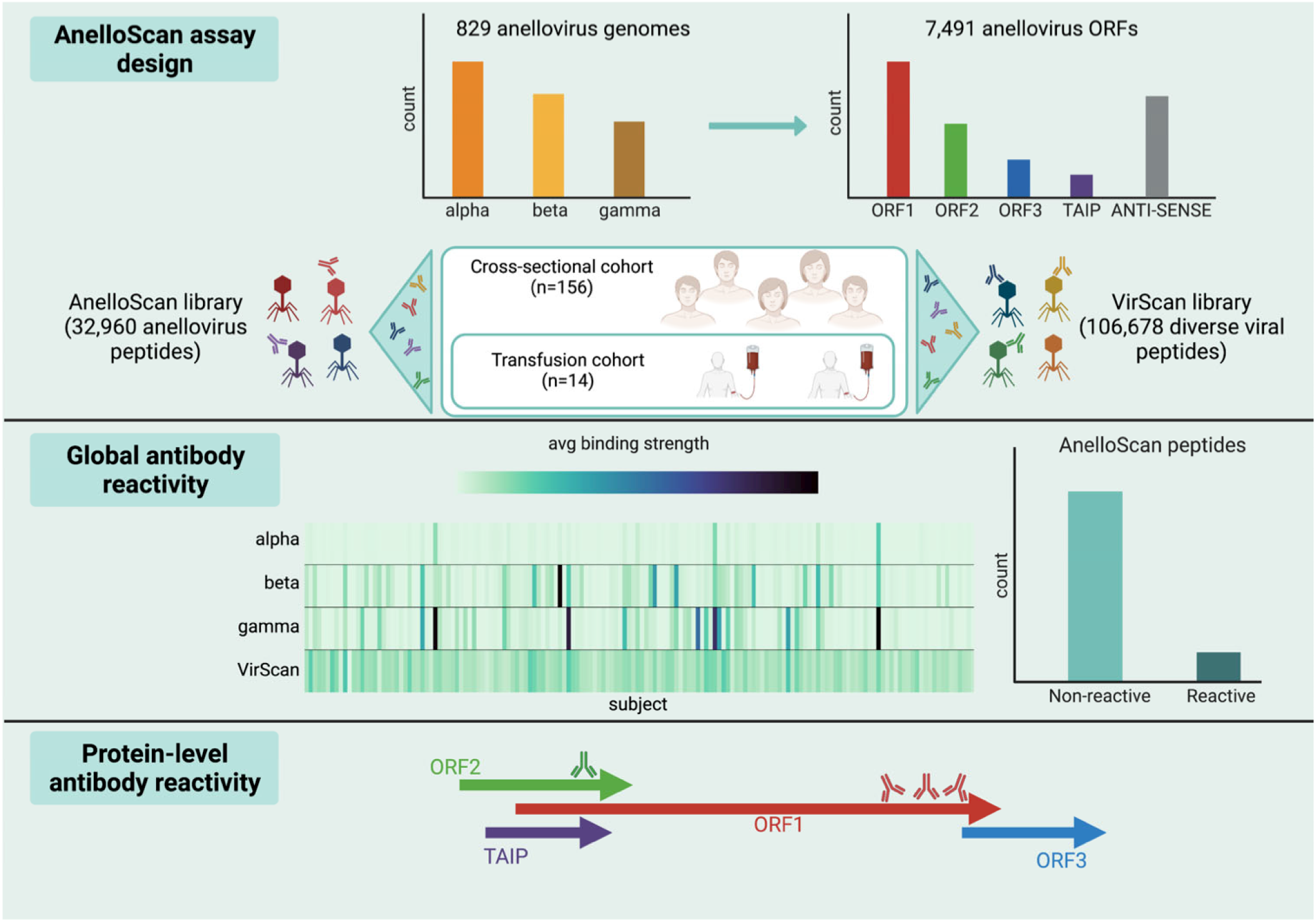

## Introduction

Commensal human viruses can establish widespread and persistent infections, and yet the nature of the host immune response to these viruses and the mechanism(s) used by the viruses to evade clearance remain unresolved. Anelloviruses make up the bulk of the viral load in healthy humans and demonstrate broad tissue tropism (De Vlaminck et al., 2013; Spandole et al., 2015). Infections with anelloviruses are ubiquitous, occur early in life and persist into adulthood, with each individual harboring a unique, personal “anellome” (Arze et al., 2021; Kaczorowska and van der Hoek, 2020). Despite increased viral load being associated with diseases such as hepatitis and cancer, no study thus far has shown evidence for their causal involvement in any disease (Kaczorowska and van der Hoek, 2020).

Anelloviruses exhibit enormous sequence heterogeneity and likely utilize recombination as a mechanism to drive their observed diversity (Arze et al., 2021; Kaczorowska and van der Hoek, 2020). They are extremely small (genome size ranging from 1.6kb to 3.9kb (Varsani et al., 2021)), non-enveloped viruses with a circular, single-stranded DNA genome that is transcribed into three distinct mRNA isoforms by means of alternative splicing (Qiu et al., 2005). The longest, singly-spliced isoform undergoes translation to give rise to two proteins. ORF1 is the longest and the putative capsid protein. ORF2 has been implicated in regulating immune response by suppressing NF-κB (Zheng et al., 2007). The two smaller, doubly-spliced mRNA isoforms code for four proteins, named ORF1/1, ORF2/2, ORF1/2 and ORF2/3, whose role in the anellovirus life cycle has yet to be elucidated. Additionally, two other ORFs have been characterized in anellovirus genomes. ORF3 is of unknown function (Arze et al., 2021), while torque teno virus TTV-derived apoptosis-inducing protein (TAIP) has been shown to induce apoptosis, preferentially in hepatocellular carcinoma cells (Kooistra et al., 2004).

Previous studies have shown that anellovirus titers are elevated in immunosuppressed individuals, suggesting that circulating anellovirus levels are under some degree of immunologic control (De Vlaminck et al., 2013). The nature of the immune response, however, is not well understood. While sero-survey studies have reported the prevalence of antibodies to anelloviruses, a broader understanding of genus- and ORF-level variations in antibody responses is needed (Gergely et al., 2005; Handa et al., 2000; Kakkola et al., 2002; Ott et al., 2000; Tsuda et al., 1999; Maggi and Bendinelli, 2009).

To characterize the immunogenicity of anelloviruses and the antigenic profile of anellovirus proteins, we employed the PhlP-Seq assay (Larman et al., 2011) using a novel “AnelloScan” T7 phage display peptide library representing close to 7,500 anellovirus open reading frame (ORF) sequences identified from DNA sequencing (DNA-seq) data. We then used the AnelloScan peptide library to screen serum samples from a cross-sectional cohort of 156 subjects for antibody binding. We observed that a majority of anellovirus peptides did not demonstrate any antibody reactivity, while the peptides that were reactive were predominantly derived from the C-terminus of ORF1. Moreover, analysis of AnelloScan data from a longitudinal blood transfusion donor(s)-recipient cohort revealed that most newly-transmitted anelloviruses did not exhibit a detectable antibody response in the recipient, and those that did exhibited a delayed response. These findings augment our understanding of host-commensal virome interactions by shedding new light on the nature of the adaptive immune response to anelloviruses.

## Results

### Design and testing of the AnelloScan phage display library

The VirScan phage display library described in a previous PhlP-Seq study consists of 106,678 peptides representing about 400 species and strains of known human viruses (Xu et al., 2015). The anellovirus component of the VirScan library comprises 420 peptides from 8 viruses (6 *Alphatorqueviruses* or “alpha”, 1 *Betatorquevirus* or “beta”, 1 *Gammatorquevirus* or “gamma”). To examine antibody reactivity against a larger and more diverse set of anellovirus sequences, we designed and constructed the “AnelloScan” phage display library representing 829 anellovirus genomes, a combination of public sequences from GenBank and sequences discovered using the AnelloScope technology to analyze 157 samples of human blood (Arze et al., 2021). The genomes included in the AnelloScan library comprise hundreds of unique sequences from the three human anellovirus genera (326 alpha, 357 beta, 146 gamma), representing an expansive increase over the anellovirus portion of the VirScan library, both in terms of number of sequences and diversity **(Fig. 1A, left)**.

**Figure 1.**
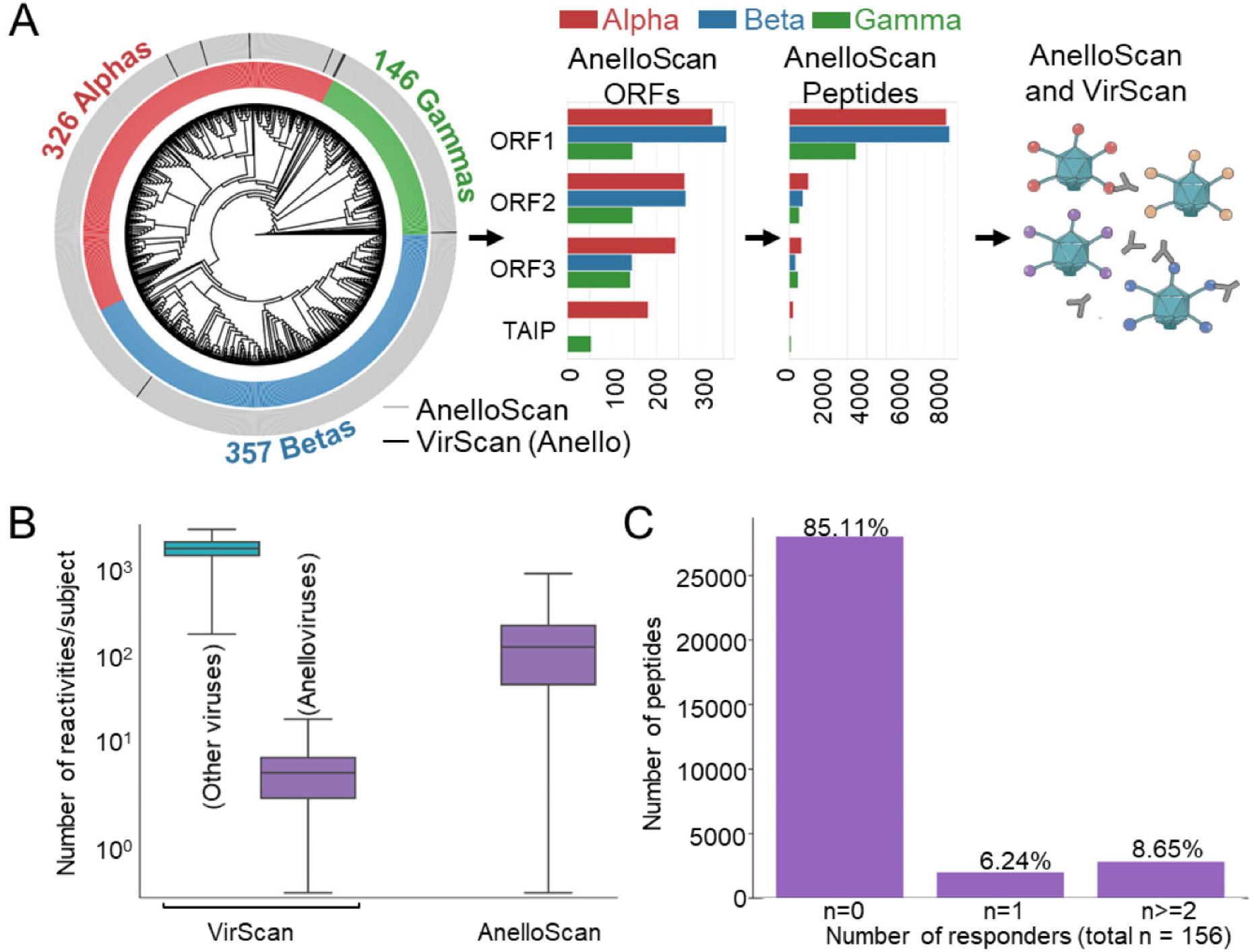
Design of the AnelloScan phage display library. **(A)** Cladogram of 837 anellovirus ORF1 sequences - 829 from the AnelloScan library (grey bars) and 8 from the VirScan library (black bars). ORFs predicted from 829 anellovirus genome sequences were classified as ORF1/ORF2/ORF3/TAIP and used to design the 56-mer AnelloScan phage display library comprising 32,960 peptides. ORFs predicted on the antisense strand were also included in the AnelloScan library **(Fig. S1).** Serum samples from a cross-sectional cohort of 156 subjects were screened against the AnelloScan and VirScan libraries. **(B)** Box plot showing the number of antibody reactivities detected in each subject against peptides in the VirScan (anellovirus – purple; all other viruses - cyan) and AnelloScan libraries. (C) Bar plot showing the number of AnelloScan peptides exhibiting an antibody reactivity in zero, one and more than one subject.

We identified ORFs on the anellovirus genome sequences using ORFfinder (National Center for Biotechnology Information, NCBI) and labelled them as ORF1/ORF2/ORF3/TAIP based on their length, location, reading frame and presence of known motifs (Methods). A total of 829 ORF1, 674 ORF2, 529 ORF3 and 234 TAIP sequences were annotated from the anellovirus genome sequences **(Fig. 1A, center, Table S1).** Additionally, ORF sequences predicted on the anti-sense strand of the anellovirus genomes, from which no known proteins are translated, were also included in the AnelloScan library **(Fig. S1).** The AnelloScan phage display library consists of 32,960 peptides, each of length 56 amino acids (aa), tiling 7,491 ORF sequences with 28 aa overlaps **(Table S1).** A cross-sectional cohort of 156 subjects were screened by both the AnelloScan and VirScan assay **(Fig. 1A, right)**.

For quality control, we included 10 public viral epitopes to serve as positive controls, as well as 10 peptides to serve as negative controls, in the AnelloScan library. None of the negative control peptides were reactive in any subject, whereas each of the positive control peptides was reactive in at least 103 out of the 156 subjects **(Fig. S2).** Additionally, to assess the reproducibility of the assay, we analyzed antibody reactivities of the AnelloScan peptides between replicate and non-replicate samples. Reactivities between replicates were highly correlated (*R^2^* = 0.791, **Fig. S3A)** while reactivities between non-replicate samples were poorly correlated (*R^2^* = 0.001, **Fig. S3B)**.

In each of the 156 subjects in the cross-sectional cohort, we measured the number of peptides in both the VirScan and AnelloScan libraries that exhibited an antibody reactivity. We found that reactivity to the 420 peptides of the anellovirus portion of the VirScan library (minimum reactivities per subject: 0; maximum: 17; median: 4) comprised a small fraction of the total reactivities observed in the rest of the 106,258 peptides in the library (minimum reactivities per subject: 171; maximum: 2920; median: 1738.5). By expanding the number of anellovirus peptides from 420 in the VirScan library to more than 32,000 in the AnelloScan library, we expected to greatly enhance the sensitivity of the assay for detecting antibody binding against anellovirus peptides. Indeed, we observed an increase in the number of anellovirus antibody reactivities detected with the AnelloScan library (minimum reactivities per subject: 0; maximum: 878; median: 127) **(Fig. 1B).** The distribution of “hitsfold change” (HFC, Methods), a measure of the strength of antibody reactivity, was significantly lower for anellovirus peptides (mean HFC = 28) compared to another chronic and highly prevalent human virus, Epstein-Barr virus (EBV, mean HFC = 35) (p=1.06×10^−49^, **Fig. S4)**.

We first assessed the number of subjects with reactivity against each peptide in the AnelloScan library. Similar to other anti-viral antibody profiles (Morgenlander et al., 2021; Shrock et al., 2020; Xu et al., 2015), the vast majority of the peptides (n=28,052; 85.11% of the library) did not display antibody reactivity in any subject tested and a further 2,058 peptides (6.24%) showed a reactivity in exactly one subject. The remaining 2,850 peptides (8.65%) displayed antibody reactivity in two or more subjects **(Fig. 1C).** Moreover, we did not detect significant differences in anellovirus antibody reactivity based on the age, gender, or race of the subject **(Fig. S5).** Taken together, these observations suggest that most anellovirus ORF-derived peptides are not associated with a detectable antibody response.

### Antibody reactivities are localized to the C-terminus of ORF1

Since the AnelloScan library included genomes from all three human genera (alpha, beta, and gamma - **Fig. 1A, left),** we next examined antibody responses by genus. We classified the peptides based on the genus of the virus from which they were derived and calculated the average HFC of peptides from each genus in all the subjects screened. We observed that antibody reactive peptides were predominantly derived from beta and gamma anelloviruses. Peptides derived from alpha anelloviruses, despite their relative overrepresentation, tended to be much less frequently reactive **(Fig. 2A)**.

**Figure 2.**
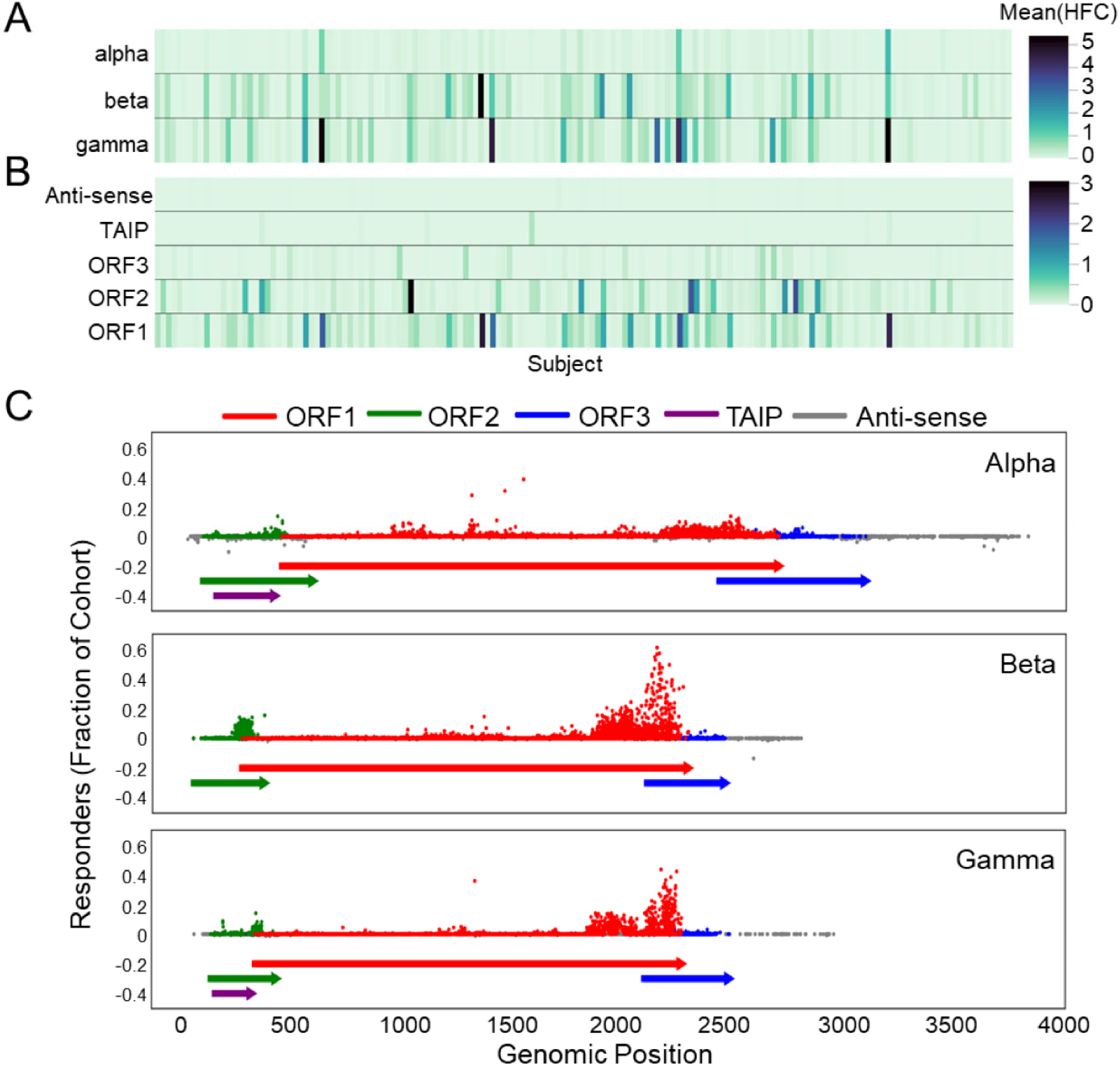
Antibody response against anellovirus peptides by genus, ORF and location. Heatmaps showing the mean hits fold change (HFC) values of peptides by **(A)** genus and **(B)** ORF. **(C)** Genome plot showing the regions of the anellovirus genomes associated with antibody reactivities. Y-axis denotes fraction of the cross-sectional cohort (n=156) responding to each peptide and x-axis the location of each peptide along the anellovirus genome.

We next calculated the average HFC of peptides from each ORF in all the subjects screened **(Fig. 2B),** revealing that the most frequently reactive peptides derived from ORF1, which is believed to be the capsid protein and a previously suspected target of immune recognition (Kaczorowska and van der Hoek, 2020). In line with findings from a previous study (Kakkola et al., 2008), we also observed less frequent reactivity to peptides from ORF2, a protein that has been shown to play a role in immune evasion (Zheng et al., 2007). In contrast, peptides derived from the other two proteins – ORF3 and TAIP – were rarely reactive. It is not currently known whether anelloviruses translate proteins from their anti-sense strand; in this study, we observed no supportive evidence for immune recognition of proteins translated from anti-sense ORFs **(Fig. 2B,** top).

Anellovirus genomes were grouped by genus and the prevalence of peptide reactivities plotted along the genome **(Fig. 2C)**. The most prevalent reactivities were associated with peptides from the C-terminal region of ORF1. Peptides representing the N-terminus and the hypervariable region in the middle of ORF1 were much less frequently reactive compared to C-terminus peptides.

### Reactive peptides localized to the ORF1 C-terminus of the three genera share a common motif

We next used a motif detection tool, Multiple Expectation-maximization for Motif Elicitation (MEME) (Bailey and Elkan, 1994; Bailey et al., 2015), to elucidate potential anellovirus antibody epitopes. Within each genus, all frequently reactive peptides (>=2.5% prevalence; alpha = 434 peptides, beta = 915 peptides, gamma = 493 peptides) were compared against all non-reactive peptides. The resulting 7, 9 and 8 motifs belonging to alpha, beta, and gamma anelloviruses, respectively, are shown in **Tables S2–S4**. Alpha anelloviruses had the least reactive motifs (range of responder fractions: 0.12-0.24), while beta anelloviruses had the most reactive motifs (range: 0.27-0.80). Gamma motif reactivity was intermediate (range: 0.21-0.63). The median HFC values for all peptides containing the motifs were comparable for all three genera. Most of the identified motifs are localized to the C-terminal portion of ORF1 across all three genera, with a smaller number of motifs localized to the ORF2 of beta and gamma anelloviruses **(Fig. 3A)**.

**Figure 3.**
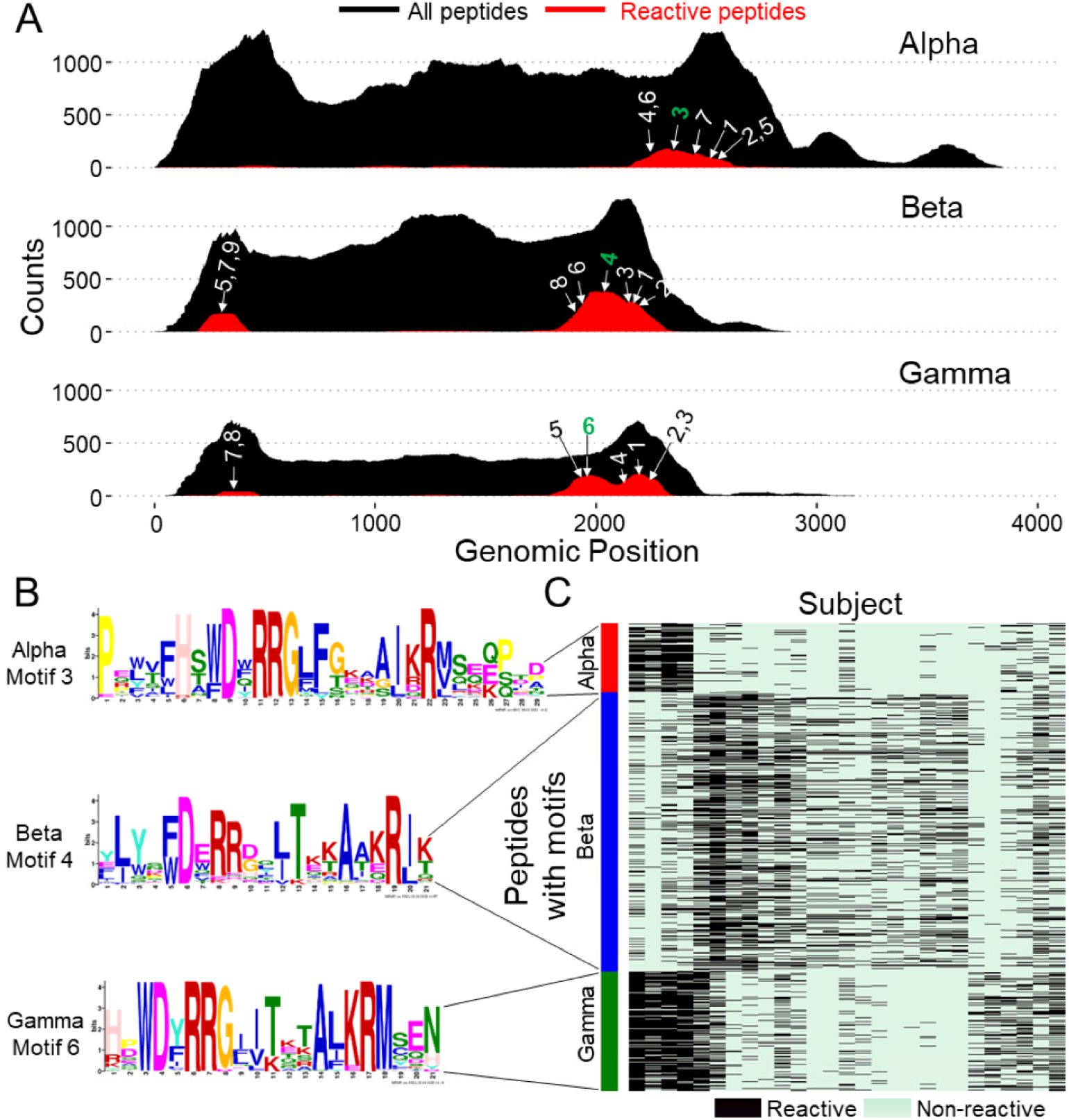
Motifs identified in reactive peptides and their cross-reactivity between anellovirus genera. **(A)** Peptide distribution map summarizing the location of the top motifs reported in **Tables S2–S4**. The area plot in black shows the depth of coverage by location of all peptides included in the library. The plot in red shows the location of peptides reactive in at least 2.5% of the cross-sectional cohort. (B) Sequence logos of the most prevalent motif shared between the three genera. **(C)** Cluster map showing antibody reactivity of peptides containing the motifs shown in B. Onlysubjects reactive against at least 10% of the epitope-containing peptides are included in the figure.

The most prevalent motif shared between the three genera is localized to the ORF1 C-terminus **(Fig. 3B)**. The fraction of the cohort reactive to peptides carrying this motif are 0.20 for alpha anelloviruses, 0.47 for beta anelloviruses and 0.31 for gamma anelloviruses **(Tables S2–S4)**. To investigate patterns of cross-reactivity among peptides carrying this motif, we clustered a heatmap of binarized reactivities **(Fig. 3C)**. Individuals with strong reactivity to peptides from alpha anelloviruses also tended to react strongly to gamma peptides, whereas there was less overlap between beta peptides and the other two genera. In summary, while peptides derived from the ORF1 C-terminus in all three genera exhibited antibody reactivity, we identified only one trans-genus epitope motif, which is likely cross-reactive between alpha and gamma anelloviruses.

### Anelloviruses can be transmitted from a donor to a recipient without generating a detectable antibody response

Since anelloviruses are transmitted from donors to recipients during blood transfusions (Arze et al., 2021), longitudinal sampling of recipients provides a potential opportunity to detect the occurrence and measure the kinetics of antibody responses to transmitted anelloviruses. The longitudinal samples used in this study consists of sera from 14 blood transfusion recipients (R01 - R14) at 5 different time points, one time point pre-transfusion (TO) and four time points post-transfusion (T1, T2, T3, T4) **(Table S2)**. By performing DNA-seq on serum samples from the recipients and their corresponding donors, we identified anelloviruses present in a donor that were transmitted to the recipient after the transfusion event (i.e., absent in recipient pre-transfusion at time point TO and present in recipient post-transfusion at one or more time points T1/T2/T3/T4). These transmitted anelloviruses from donors to 5 recipients were included as part of the AnelloScan library. In these 5 recipients we examined antibody reactivities targeting the newly transmitted donor anelloviruses.

In three recipients (R07, R10 and R11), none of the five transmitted donor anelloviruses was associated with a post-transfusion antibody response. In the remaining two recipients, 12 of 19 (in recipient R04) and 12 of 16 (in recipient R05) transmitted anelloviruses were not associated with a post-transfusion antibody response. Of the remaining seven and four transmitted donor anelloviruses that were associated with an antibody response, the reactivities were typically not detected at the first time point post-transfusion (56 and 24 days post-transfusion for R04 and R05, respectively). Instead, in all but one of these 11 cases, antibody reactivities against transmitted anelloviruses were first detected −150 and −100 days post-transfusion for R04 and R05, respectively **(Fig. 4)**. This response represents relatively delayed kinetics compared with the conventional seroconversion associated with pathogenic viral infections (~14 days after infection) (Cohen et al., 2011). Moreover, there were multiple instances where we detected antibody reactivity to transmitted anelloviruses only in the recipient and not in any of the donors. These observations suggest that the antibodies recognizing transmitted anelloviruses were not passively transmitted from a donor to the recipient during transfusion. Rather, these antibody responses were likely elicited by a newly-acquired anellovirus infection.

**Figure 4.**
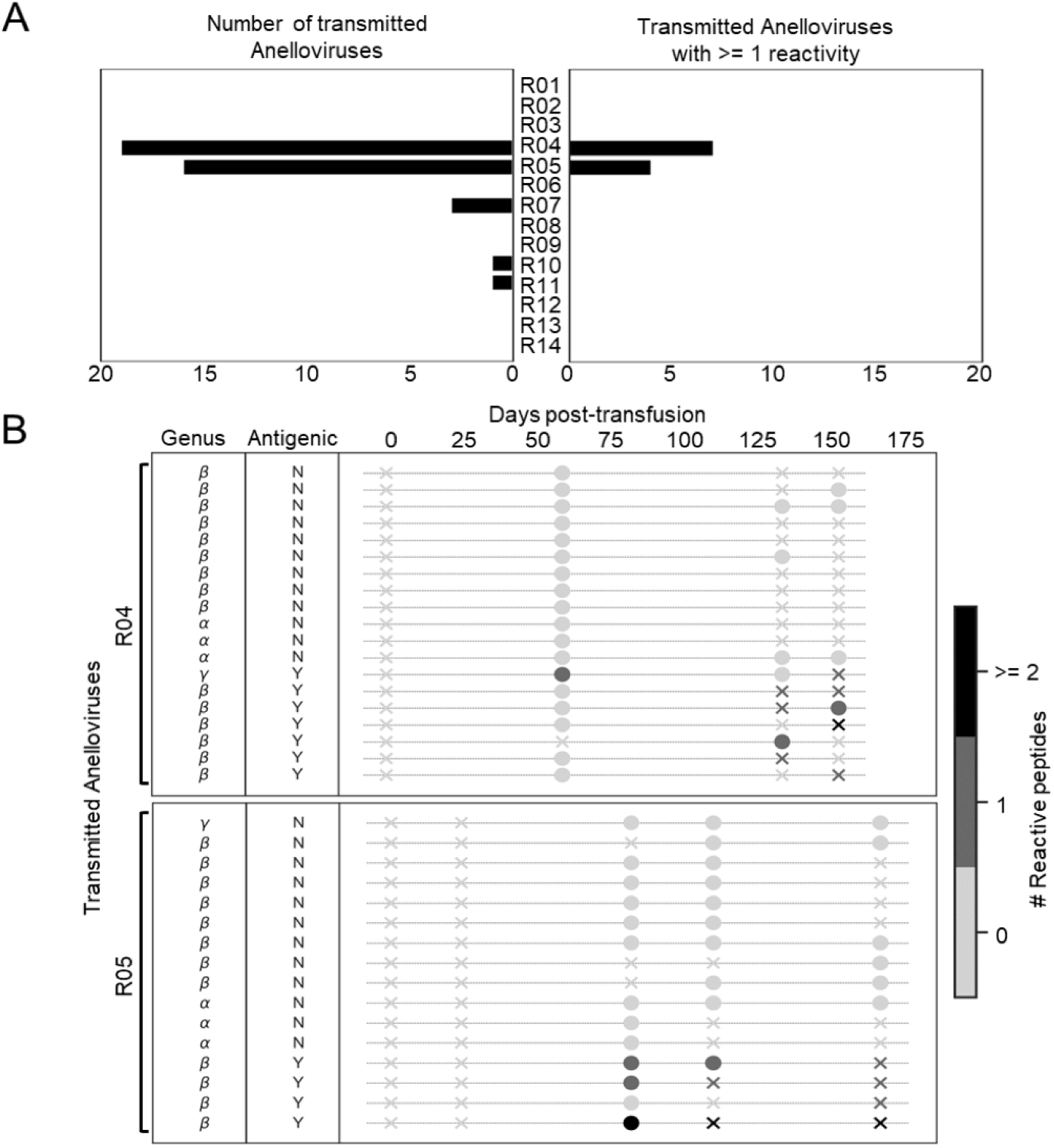
Antibody reactivities against transmitted anelloviruses in blood transfusion recipients. **(A)** Bar plot showing the number of transmitted anelloviruses in 14 blood transfusion recipients. The left panel shows the number of viruses transmitted from a donor to each recipient, as measured by DNA-seq, and the right panel shows the number of such viruses that evoked an immune response (defined as reactivity to at least one peptide from a virus) in the recipient post-transfusion, as measured by AnelloScan. (B) Antibody reactivities against transmitted anelloviruses in blood transfusion recipients R04 and R05 using AnelloScan. “antigenic”: “N” - virus did not generate an antibody response, “Y” - virus generated an antibody response. X’s and O’s indicate absence and presence of the virus based on DNA-seq data, respectively, shaded according to number of reactive peptides.

We further assessed the longitudinal stability of anti-anellovirus antibody reactivities in the 14 blood transfusion recipients by calculating the correlation of all AnelloScan peptides’ HFC values between different time points within a recipient as well as between recipients. While inter-individual variability is extremely high, as observed in the cross-sectional cohort, the intra-individual reactivity profile remained relatively stable over time **(Fig. S6A)**. Furthermore, AnelloScan reactivity profiles in donors were completely distinct from the recipients; no significant correlations in reactivity were observed in the early post-transfusion time points **(Fig. S6B),** further indicating that passive transfer of antibodies from donor to recipient did not confound this analysis.

Finally, we asked if there was any correlation between the detection of specific anellovirus genomes and the detection of antibody reactivity by AnelloScan. AnelloScope and AnelloScan were both performed on a set of 58 serum samples from the blood transfusion cohort (Arze et al., 2021). We mapped DNA-seq reads from each sample to the 829 anellovirus genome sequences in the AnelloScan library and tested for an association between genomes and antibody reactivity. We detected no significant correlation between the detection of a viral genome and the presence of reactivity against that virus (Fisher’s exact test; p-value: 0.58).

## Discussion

While anelloviruses chronically infect most humans, higher viral titers have been reported in immunocompromised individuals including solid organ transplant recipients as well as patients with HIV or cancer (Doberer et al., 2020; Uhl et al., 2021). These observations suggest that anellovirus levels are controlled by a healthy immune system. Previous studies have shown that circulating antibodies are recognized by, and form complexes with anelloviruses, suggesting that anellovirus-specific immune responses are, at least in part, humoral in nature (Mankotia and Irshad, 2014). Correspondingly, detection of anti-anellovirus antibodies using the C-terminal fragment of the ORF1 protein has been reported by multiple groups (Maggi and Bendinelli, 2009). ORF1 is considered the putative anellovirus capsid protein and therefore a natural target for immune recognition (Kaczorowska and van der Hoek, 2020). In limited studies, both ORF1 and ORF2 from *Alphatorqueviruses* were found to be recognized by antibodies in the serum of anellovirus-infected individuals (Gergely et al., 2005; Kakkola et al., 2008). Our findings in this study are largely concordant with these observations and extend them by simultaneously characterizing anti-anellovirus antibody specificities at the genus and ORF level, as well as identifying potential cross-reactivities.

The ORF1 protein is translated from the larger singly-spliced mRNA isoform, whereas the smaller ORF1/1 protein is translated from one of the two smaller doubly-spliced mRNA isoforms, producing identical C-terminal fragments. The prevalent antibody response against the C-terminus of ORF1 cannot be specifically assigned to either one of these isoforms.

The genomes of anelloviruses are negative-sense, single-stranded DNA. Prior to transcription, the genome is converted to double-stranded DNA in the nucleus of the infected cell (Shulman and Davidson, 2017). It is unknown whether anelloviruses produce non-canonical proteins from mRNA transcribed from their negative-sense DNA. We represented these potential viral protein products in the AnelloScan library, but did not observe a measurable reactivity to these proteins.

While we observed multiple cases of anelloviruses being transmitted from a donor to a recipient without eliciting a detectable antibody response, this observation could be an artifact of the time points at which the recipient serum samples were collected – ranging from less than a week to more than three months for the first time point after the transfusion event. A higher resolution longitudinal study design would enhance our understanding of the kinetics of the antibody response to newly-acquired anellovirus infections.

The commensalism and potential immune evasion of anelloviruses nominate them as candidates for delivery of therapeutic payloads (Arze et al., 2021; Sandbrink et al., 2022). To date, immune responses to recombinant adeno-associated virus (AAV) vectors have been a major obstacle (Verdera et al., 2020), underscoring the need for new vector candidates. Our findings suggest that anellovirus-based - particularly alpha anellovirus-based – gene therapies have the potential for multi-dose regimens.

While the PhlP-Seq assay enables comprehensive antibody binding analysis, it is important to note that phage display libraries such as AnelloScan do not present highly conformational, discontinuous, or post-translationally modified epitopes (Mohan et al., 2018). This study may therefore suffer from a high rate of false negative detection (low sensitivity).

In summary, PhlP-seq using the novel AnelloScan phage display library has enabled the first comprehensive, unbiased, cohort-scale characterization of the antibody response to the commensal human anellome and opens the door for further investigation into the roles of anelloviruses in health and disease.

## Methods

### Curation of anellovirus genomes

Anellovirus genome sequences were collated and filtered from both internal sequencing projects and publicly available complete anellovirus genomes from the NCBI GenBank repository. Sequences were size selected between 2,000bp and 4,000bp utilizing the seqkit software (Shen et al., 2016) with the parameters −m 2000 and −M 4000. Sequences were re-oriented to an arbitrary start position of the 5’ UTR using the nhmmer software (Wheeler and Eddy, 2013). A set of 829 anellovirus genome sequences were included in the AnelloScan library.

### Identification of ORFs

To predict coding sequences for peptide synthesis, ORFs were called on the 829 anellovirus genomes in the AnelloScan library using ORFfinder (NCBI) with the following parameters: *-strand both −s 1 −outfmt 1 −ml 150*.

### Annotation of ORFs

The ORFs predicted by ORFfinder were labelled as ORF1IORF2/0RF3/TAIP based on a linear workflow employing the criteria outlined below:

1. An ORF was labelled as ORF1 if:

a. The length of the ORF was >= 500 aa
2. An ORF was labelled as ORF2 if:

a. The length of the ORF was >= 75 aa
b. The ORF spanned the start of ORF1
c. The ORF was not from the same reading frame as ORF1
d. The ORF contained the conserved motif in ORF2 (Takahashi et al., 2000)
3. An ORF was labelled as ORF3 if:

a. The length of the ORF was >= 50 aa
b. The ORF spanned the end of ORF1
c. The ORF was not from the same reading frame as ORF1 or ORF2
d. The ORF contained one of the two conserved motifs in ORF3 (Arze et al., 2021)
4. An ORF was labelled as TAIP if:

a. The length of the ORF was >= 50 aa
b. The ORF spanned the start of ORF1 and ORF2
c. The ORF was not from the same reading frame as ORF1 or ORF2

### Alignment of peptides with ORFs

Since some regions of anellovirus proteins are highly conserved, a peptide might represent ORFs from more than one virus, which could in turn result in cross-reactivity. Therefore, we performed a BLAST alignment of all the AnelloScan peptides against all AnelloScan ORF sequences to assign each peptide to one or more ORFs. An ORF was assigned to a peptide if at least 95% of the peptide’s residues had a positive-scoring match to the ORF and the *e-value* of the BLAST hit was less than 1.

### Construction of the ORF1 phylogenetic tree

837 ORF1 amino acid sequences - 829 from the AnelloScan library and 8 from the VirScan library - were aligned using MAFFT with the G-INS-i setting (Katoh et al., 2002). The aligned ORF1 sequences from MAFFT were used to construct a maximum likelihood phylogenetic tree with RAxML (CAT sequence evolution model, BLOSUM62 substitution matrix) (Stamatakis, 2014).

### AnelloScan library design

A more detailed version of the library design process is provided in (Mohan et al., 2018). In brief, the AnelloScan library was designed from the selected ORFs using the pepsyn Python package (https://github.com/lasersonlab/pepsyn). Pepsyn (i) split the protein sequences into peptide tiles (56 aa long, 28 aa overlap), (ii) selected representative peptide sequences from peptide clusters of similarity greater than 95% (using the cd-hit python package), (iii) reverse-translated the selected peptide tiles with an optimized *E. coli* codon usage algorithm, (iv) added forward and reverse PCR primer binding sequences to the resulting DNA sequences, and (v) removed *EcoRI* and *Hindlll* restriction cloning sites by silent codon substitution. The designed library consisted of 32,960 peptides representing all three human anellovirus genera.

### AnelloScan library synthesis and cloning

Oligonucleotide library synthesis was performed by Twist Bioscience (San Francisco, CA). Mid-copy T7 phage vector (T7FNS2) derived from T7Select 10-3b (EMD Millipore, Cat No. 70548) was used for cloning. Vector DNA was digested overnight with *EcoRI/Hindlll* restriction enzymes (NEB Cat Nos. R3101, R3104) followed by a 10-minute phosphatase treatment (Quick CIP, NEB Cat No. M0525) to dephosphorylate the ends. Vector arms were gel-purified in 1% SeaPlaque agarose (Lonza, Cat No. 50105) in 1X TAE buffer.

The oligo library was dissolved in molecular biology-grade water to 100 nM. PCR amplification was performed for 12 cycles to add adapters and generate adequate amounts of insert for cloning. PCR-amplified product was column-purified (Nucleospin column, Machery-Nagel, Cat No. 740609), digested with *EcoRI/Hindlll* and gel-purified.

5 μl ligation reactions were set up with a total of 500 ng DNA (vector and insert at a 1:4 molar ratio) and high-concentration T4 DNA ligase (NEB Cat No. M0202T). The ligation mix was packaged using the T7Select Packaging Extract (EMD Millipore, Cat No. 70014). An adequate number of ligation/packaging reactions were set up to obtain 100 times as many plaques as members of the library. 20 plaques from the AnelloScan library were picked individually and their inserts amplified by PCR. Amplified inserts were Sanger sequenced to identify the inserts and evaluate them for the presence of mutations. 18 (90%) of the 20 inserts showed no mutations and two inserts showed either a non-synonymous mutation or a single base pair deletion.

### Phage ImmunoPrecipitation Sequencing

The standard protein A/G PhlP-Seq protocol, used in this study, has been previously described in detail (Mohan et al., 2018). Briefly, an enzyme-linked immunosorbent assay (ELISA) was performed to measure total IgG in serum samples and input volume was adjusted to 2 μg of IgG input per immunoprecipitation (IP). The AnelloScan and the VirScan libraries were mixed with diluted serum at 10^5^-fold library coverage. The mixture was allowed to rotate overnight at 4°C, followed by a 4-hour IP with protein A- and protein G-coated magnetic beads (Dynabeads, Invitrogen). A first PCR was performed to amplify the peptide inserts. A second PCR added adapters and indices for Iliumina sequencing. The FASTQ sequencing file was demultiplexed and aligned to obtain read count values for each peptide in the library, followed by the detection of antibody reactivity by comparison with a set of mock IPs run on the same plate, using the Phipmake R package (Sie, 2020).

For a peptide to be considered reactive, the read count needed to exceed 15, the fold change needed to be at least 5 and the p-value of differential abundance needed to be lower than 0.001 (fold-changes and p-values were calculated using the EdgeR software) (McCarthy et al., 2012; Robinson et al., 2010). Fold change values that fulfilled these three criteria were referred to as hits fold change (HFC) and the HFC of any peptide that failed at least one criterion was set to 1 (unenriched with respect to mock).

### DNA-seq of longitudinal blood transfusion donor(s)-recipient cohort

Nucleic acids were extracted from the serum with a PureLink viral DNA/RNA kit from Invitrogen. The samples were processed according to the manufacturer’s protocol with an increase to 60 minutes for the Proteinase K incubation step. Samples were eluted in 50 μL of nuclease-free water. The extracted DNA was amplified using a rolling circle amplification (RCA) protocol as previously described (Arze et al., 2021). Anellovirus genomes in the samples were amplified with the pan-anello primers developed by (Ninomiya et al., 2008). 2 μL of sample was added to 1x PCR Master Mix (Sigma-Aldrich, Basel, Switzerland) and the 4 degenerate primers at a final concentration of 1 μM each in a final volume of 25 μL. Positive samples were identified by the presence of the 128-base pair band in a 2% agarose gel. Post-RCA DNA was diluted to a volume of 50 μL to reduce viscosity of the samples and the DNA concentration was assessed by Qubit. Post-RCA DNA was prepared for sequencing using the Nextera DNA kit (Illumina, San Diego, USA). The samples were prepared following the manufacturer’s protocol for 100-500 ng input. Library QC was carried out with the D5000 screen tape on an Agilent Tapestation 4200. All libraries were then sequenced on an Illumina NextSeq 550.

Raw DNA sequence base calls were processed from multiple Illumina iSeq and NextSeq runs to produce the representative set of genomes used as references for AnelloScan. Base calls were demultiplexed using the picard suite’s ExtractllluminaBarcodes, llluminaBasecalls ToSam and MergeSamFiles utilities (Broad Institute, 2018) to produce unmapped BAM files containing short-read sequences. BAM files were further processed using the SamToFastq utility in picard to produce paired-end FASTQ files that served as a base for all downstream processing and analysis.

Paired end FASTQ files were subjected to a sequence of quality control and decontamination steps to remove adapters, lab contaminants and host background sequences. First, low quality sequences and adapter sequences were removed with the bbduk tool (Bushnell, 2014) using the following parameters: *ktrim*=*r*, *k*=*23*, *mink*=*11*, *tpe*=*t*, *tbo*=*t*, *qtrim*=*rl*, *trimq*=*20*, *minlength*=*50*, *maxns*=*2*. The adapter and contaminant reference file used in this step was created by pulling common adapter sequences from NCBI’s GenBank repository.

Next, human sequence decontamination was conducted in two passes using both the NextGenMap (Sedlazeck et al., 2013) and BWA tools (Li and Durbin, 2009). NextGenMap was executed using the following parameters, *--affine, −s 0.7*, and *-p* and BWA was executed using the *mem* algorithm with default parameters. The hg19 human reference was used in both decontamination steps.

Both rRNA and common bacterial lab contaminants were removed using the bbmap tool (Bushnell, 2014) with the following parameters: *minid*=*0.95*, *bwr*=*0.16*, *bw*=*12*, *quickmatch*=*t*, *fast*=*t*, *minhits*=*2*.

Cleaned and decontaminated reads were subjected to a deduplication step using the clumpify tool (Bushnell, 2014) with the *dedupe*=*t* parameter. Finally, the cleaned reads were mapped to the 829 AnelloScan genome sequences with bwa using the *mem* algorithm (Li and Durbin, 2009). Alignments with an alignment score (AS) less than 140 were filtered out.

## Author contributions

Conceptualization: HBL, NLY, VM; Sample collection, processing, and data generation: TV, SMJ, AB, TD; Data analysis: TV, HS, CAA; Writing: TV, HS, CAA, AB, DMN, SD, NLY, HBL.

## Competing Interest Statement

TV declares no competing interests. HS, CAA, SMJ, AB, TD, DMN, SD, and NLY are employees of and hold equity interests in Ring Therapeutics. VM is an employee of and holds equity interests in Flagship Pioneering, which also holds an equity interest in Ring Therapeutics. HBL is an inventor on an issued patent (US20160320406A) filed by Brigham and Women’s Hospital that covers the use of the VirScan technology, is a founder of ImmunelD, Portal Bioscience and Alchemab, and is an advisor to TScan Therapeutics.

## Acknowledgements

We thank Kristian G. Andersen and Gytis Dudas for assistance with data analysis and visualization. This study utilized TTVS research materials obtained from the NHLBI Biological Specimen and Data Repository Information Coordinating Center and does not necessarily reflect the opinions or views of the TTVS or the NHLBI.

## Funding

This study was funded by Ring Therapeutics.

## Supplementary Material

**Table S1.**
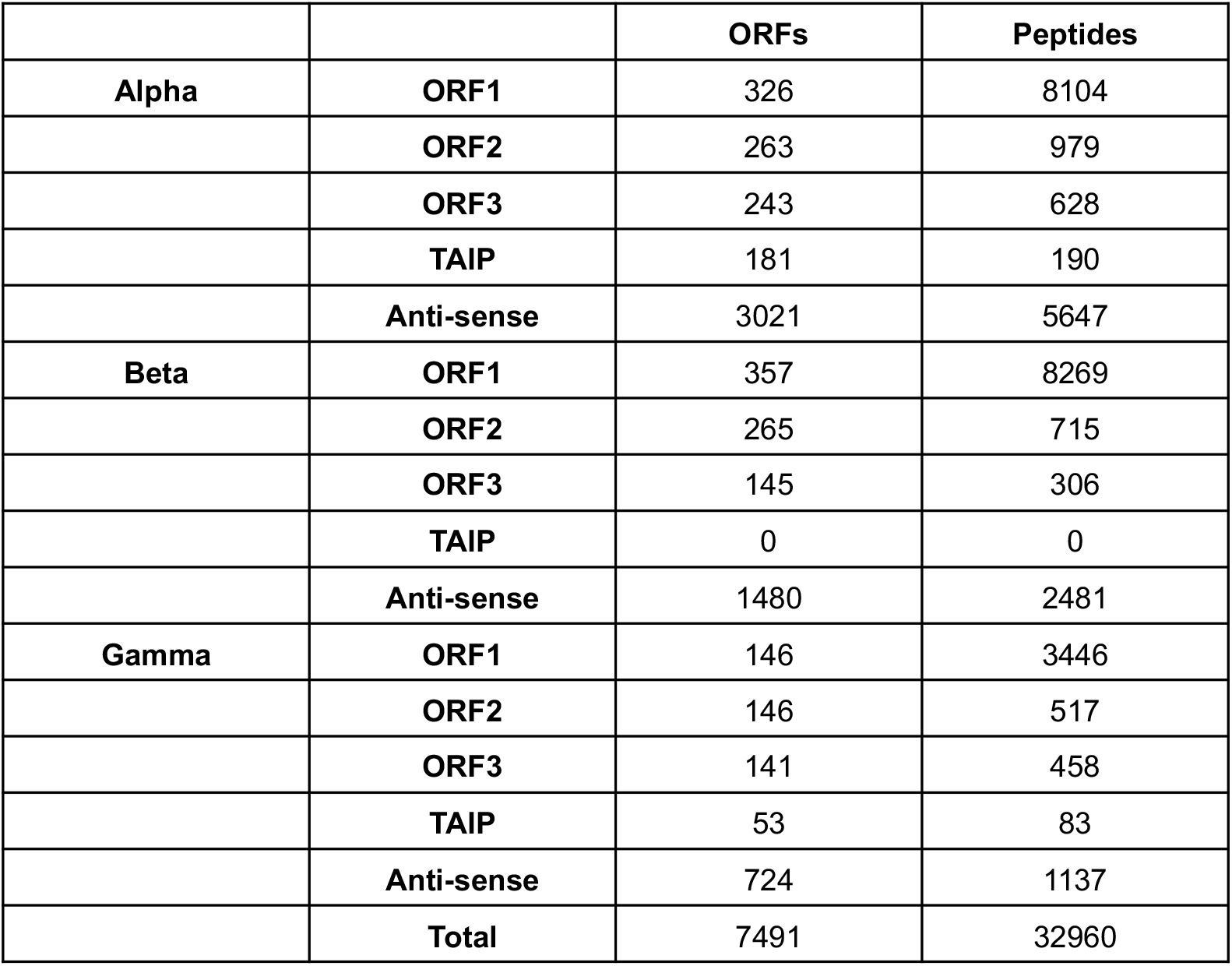
Composition of the AnelloScan library by anellovirus genus and ORF.

|  |  | ORFs | Peptides |
| --- | --- | --- | --- |
| **Alpha** | **ORF1** | 326 | 8104 |
|  | **ORF2** | 263 | 979 |
|  | **ORF3** | 243 | 628 |
|  | **TAIP** | 181 | 190 |
|  | **Anti-sense** | 3021 | 5647 |
| **Beta** | **ORF1** | 357 | 8269 |
|  | **ORF2** | 265 | 715 |
|  | **ORF3** | 145 | 306 |
|  | **TAIP** | 0 | 0 |
|  | **Anti-sense** | 1480 | 2481 |
| **Gamma** | **ORF1** | 146 | 3446 |
|  | **ORF2** | 146 | 517 |
|  | **ORF3** | 141 | 458 |
|  | **TAIP** | 53 | 83 |
|  | **Anti-sense** | 724 | 1137 |
|  | **Total** | 7491 | 32960 |

**Table S2.**
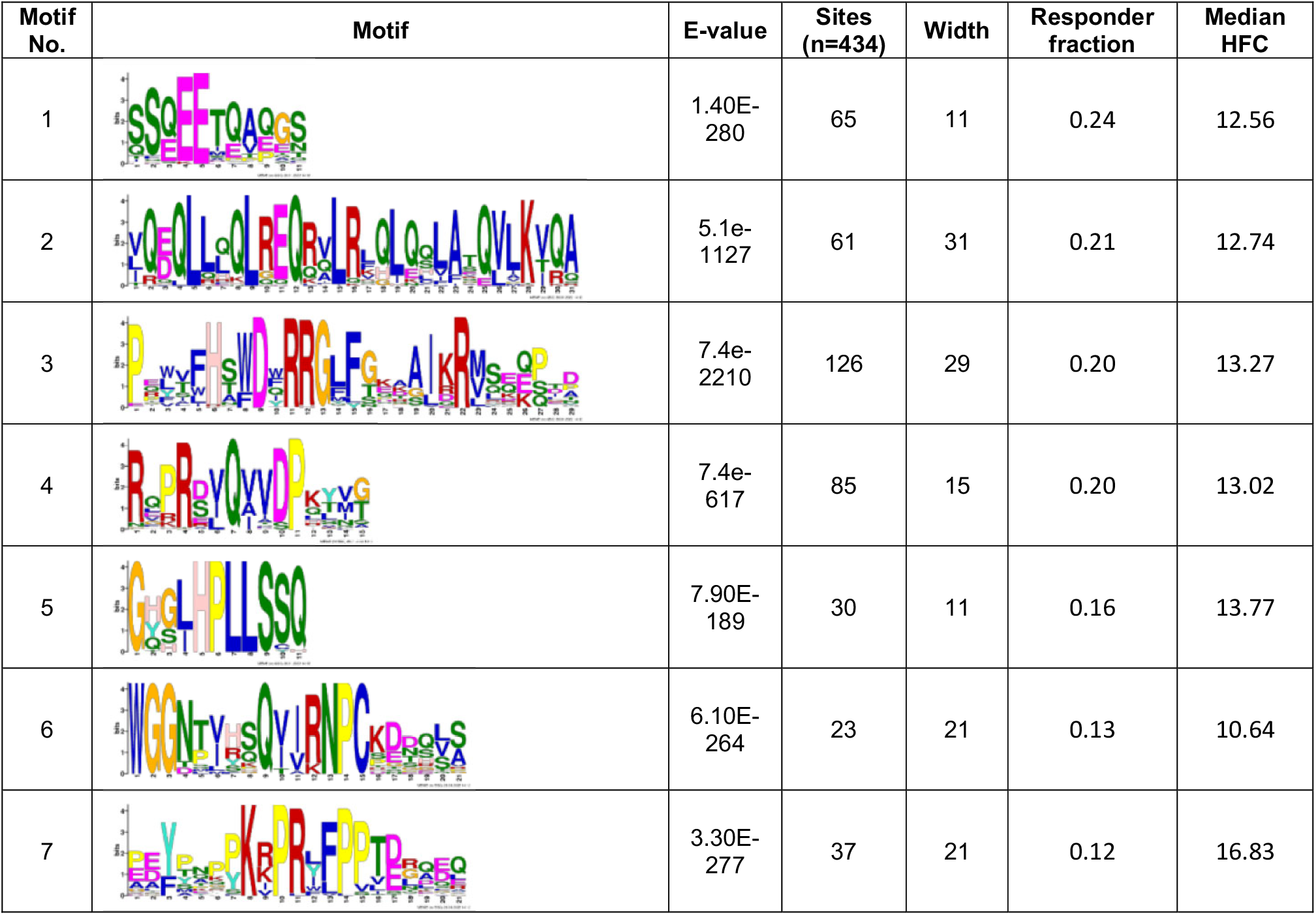
Motifs identified in reactive peptides of alpha anelloviruses. Table shows the top 7 motifs identified in alpha anellovirus peptides that are reactive in at least 2.5% of the cross-sectional cohort. A total of 434 reactive peptides were selected for analysis. E-values, number of sites containing the motif and the width of the motif reported by MEME are shown in the table above. The fraction of the cross-sectional cohort that responds to at least 3 peptides containing the motif is shown as “Responder fraction” and the median HFC of all the motif-containing reactive peptides is shown as “Median HFC”.

| Motif No. | Motif | E-value | Sites (n=434) | Width | Responder fraction | Median HFC |
| --- | --- | --- | --- | --- | --- | --- |
| 1 |  | 1.40E-280 | 65 | 11 | 0.24 | 12.56 |
| 2 |  | 5.1e-1127 | 61 | 31 | 0.21 | 12.74 |
| 3 |  | 7.4e-2210 | 126 | 29 | 0.20 | 13.27 |
| 4 |  | 7.4e-617 | 85 | 15 | 0.20 | 13.02 |
| 5 |  | 7.90E-189 | 30 | 11 | 0.16 | 13.77 |
| 6 |  | 6.10E-264 | 23 | 21 | 0.13 | 10.64 |
| 7 |  | 3.30E-277 | 37 | 21 | 0.12 | 16.83 |

**Table S3.**
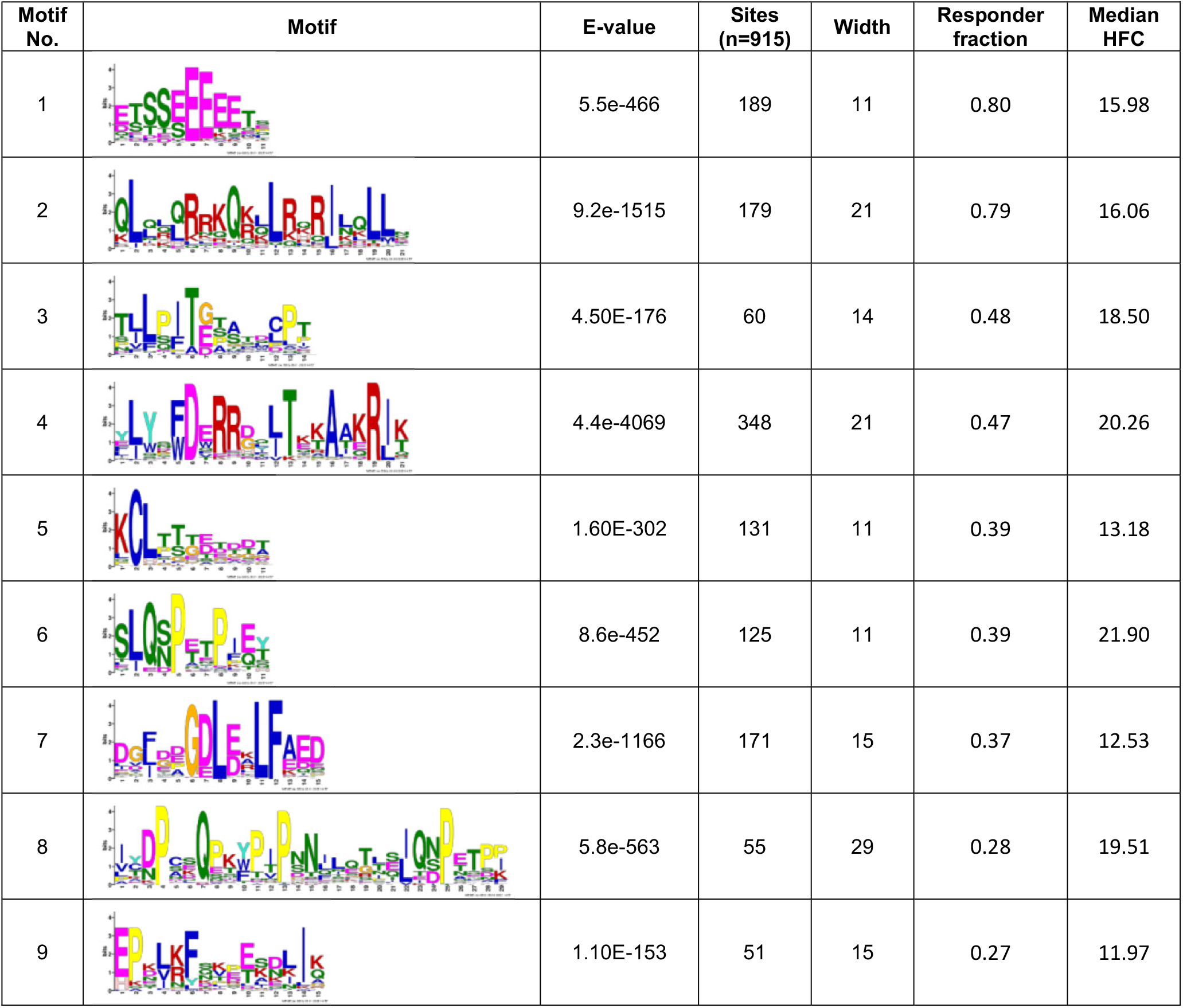
Motifs identified in reactive peptides of beta anelloviruses. Table shows the top 9 motifs identified in beta anellovirus peptides that are reactive in at least 2.5% of the cross-sectional cohort. A total of 915 reactive peptides were selected for analysis.

| Motif No. | Motif | E-value | Sites (n=915) | Width | Responder fraction | Median HFC |
| --- | --- | --- | --- | --- | --- | --- |
| 1 |  | 5.5e-466 | 189 | 11 | 0.80 | 15.98 |
| 2 |  | 9.2e-1515 | 179 | 21 | 0.79 | 16.06 |
| 3 |  | 4.50E-176 | 60 | 14 | 0.48 | 18.50 |
| 4 |  | 4.4e-4069 | 348 | 21 | 0.47 | 20.26 |
| 5 |  | 1.60E-302 | 131 | 11 | 0.39 | 13.18 |
| 6 |  | 8.6e-452 | 125 | 11 | 0.39 | 21.90 |
| 7 |  | 2.3e-1166 | 171 | 15 | 0.37 | 12.53 |
| 8 |  | 5.8e-563 | 55 | 29 | 0.28 | 19.51 |
| 9 |  | 1.10E-153 | 51 | 15 | 0.27 | 11.97 |

**Table S4.**
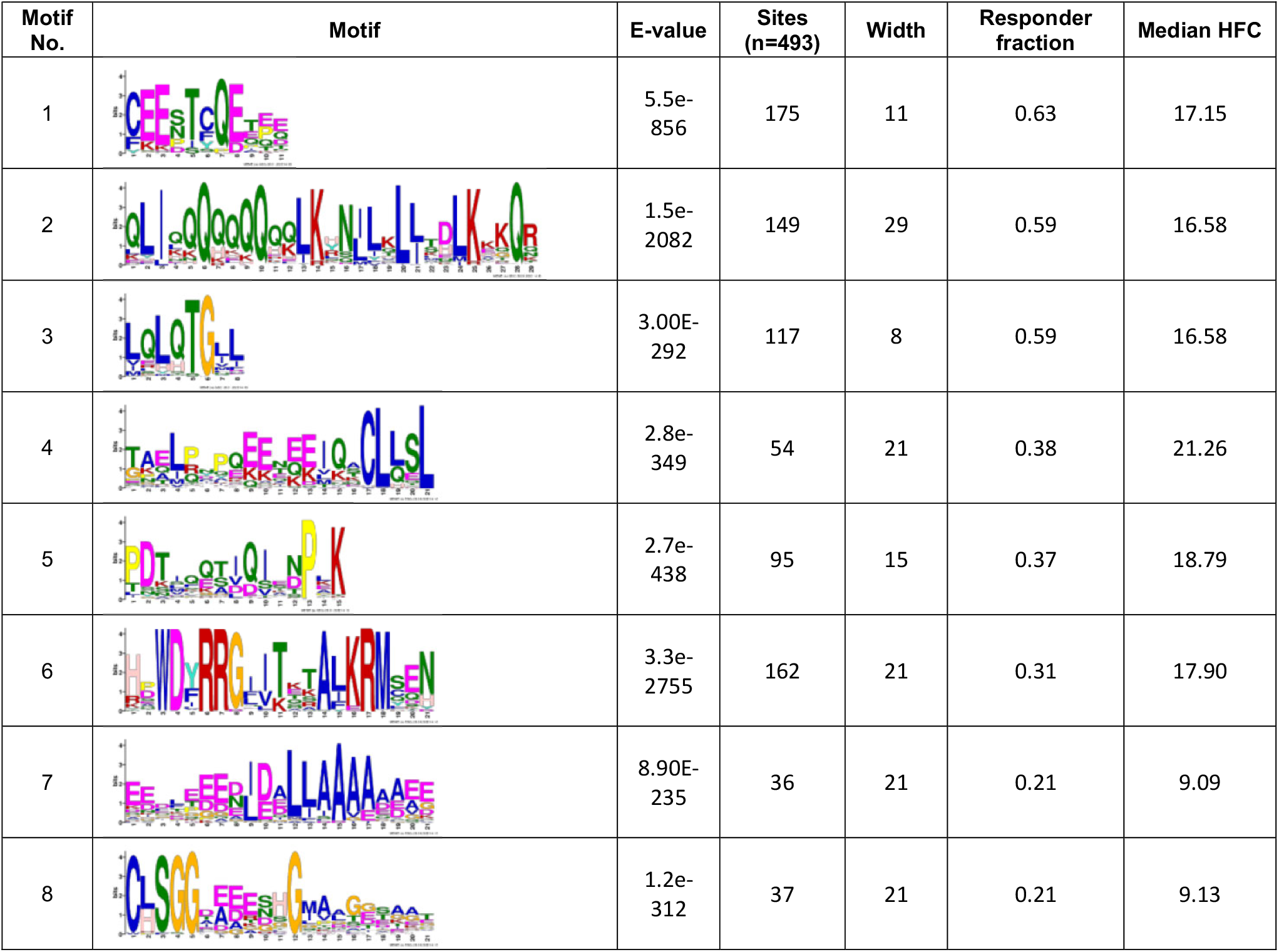
Motifs identified in reactive peptides of gamma anelloviruses. Table shows the top 8 motifs identified in gamma anellovirus peptides that are reactive in at least 2.5% of the cross-sectional cohort. A total of 493 reactive peptides were selected for analysis.

| Motif No. | Motif | E-value | Sites (n=493) | Width | Responder fraction | Median HFC |
| --- | --- | --- | --- | --- | --- | --- |
| 1 |  | 5.5e-856 | 175 | 11 | 0.63 | 17.15 |
| 2 |  | 1.5e-2082 | 149 | 29 | 0.59 | 16.58 |
| 3 |  | 3.00E-292 | 117 | 8 | 0.59 | 16.58 |
| 4 |  | 2.8e-349 | 54 | 21 | 0.38 | 21.26 |
| 5 |  | 2.7e-438 | 95 | 15 | 0.37 | 18.79 |
| 6 |  | 3.3e-2755 | 162 | 21 | 0.31 | 17.90 |
| 7 |  | 8.90E-235 | 36 | 21 | 0.21 | 9.09 |
| 8 |  | 1.2e-312 | 37 | 21 | 0.21 | 9.13 |

**Table S5.** Days post-transfusion for the longitudinal blood transfusion recipient cohort. Samples were collected at one time point pre-transfusion (TO) and four time points post-transfusion (T1 to T4) from 14 transfusion recipients (R01 to R14). The values indicate the number of days relative to transfusion. X indicates a missing sample.

| Sample | T0 | T1 | T2 | T3 | T4 |
| --- | --- | --- | --- | --- | --- |
| R01 | -1 | 82 | 105 | 132 | 154 |
| R02 | -1 | 7 | 71 | 106 | 166 |
| R03 | -8 | 41 | 70 | 127 | 176 |
| R04 | -1 | 56 | 126 | 144 | X |
| R05 | -1 | 24 | 82 | 110 | 167 |
| R06 | -1 | 7 | 19 | 32 | 59 |
| R07 | -1 | 28 | 55 | 107 | 167 |
| R08 | -1 | 11 | 59 | 86 | 108 |
| R09 | -1 | 14 | 57 | 107 | 125 |
| R10 | -1 | 6 | 18 | 71 | 150 |
| R11 | -2 | 44 | 129 | 148 | 269 |
| R12 | 0 | 5 | 44 | 52 | 67 |
| R13 | -1 | 6 | 42 | 56 | 84 |
| R14 | -4 | 97 | 146 | 167 | 275 |

**Figure S1.**
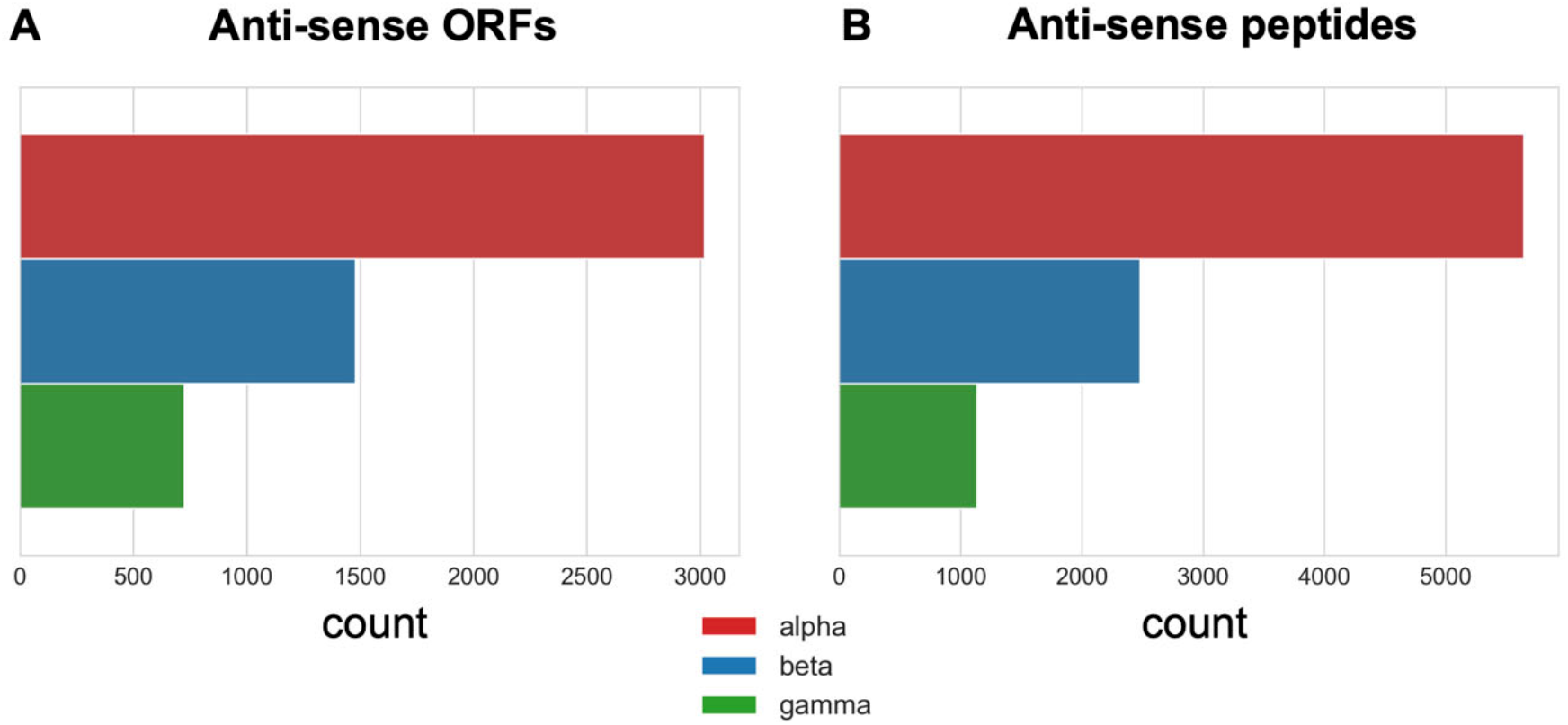
Sense and antisense library ORFs and peptides. Counts of **(A)** anti-sense ORFs (n=5,225) and **(B)** anti-sense peptides (n=9,265) in the AnelloScan library.

**Figure S2.**
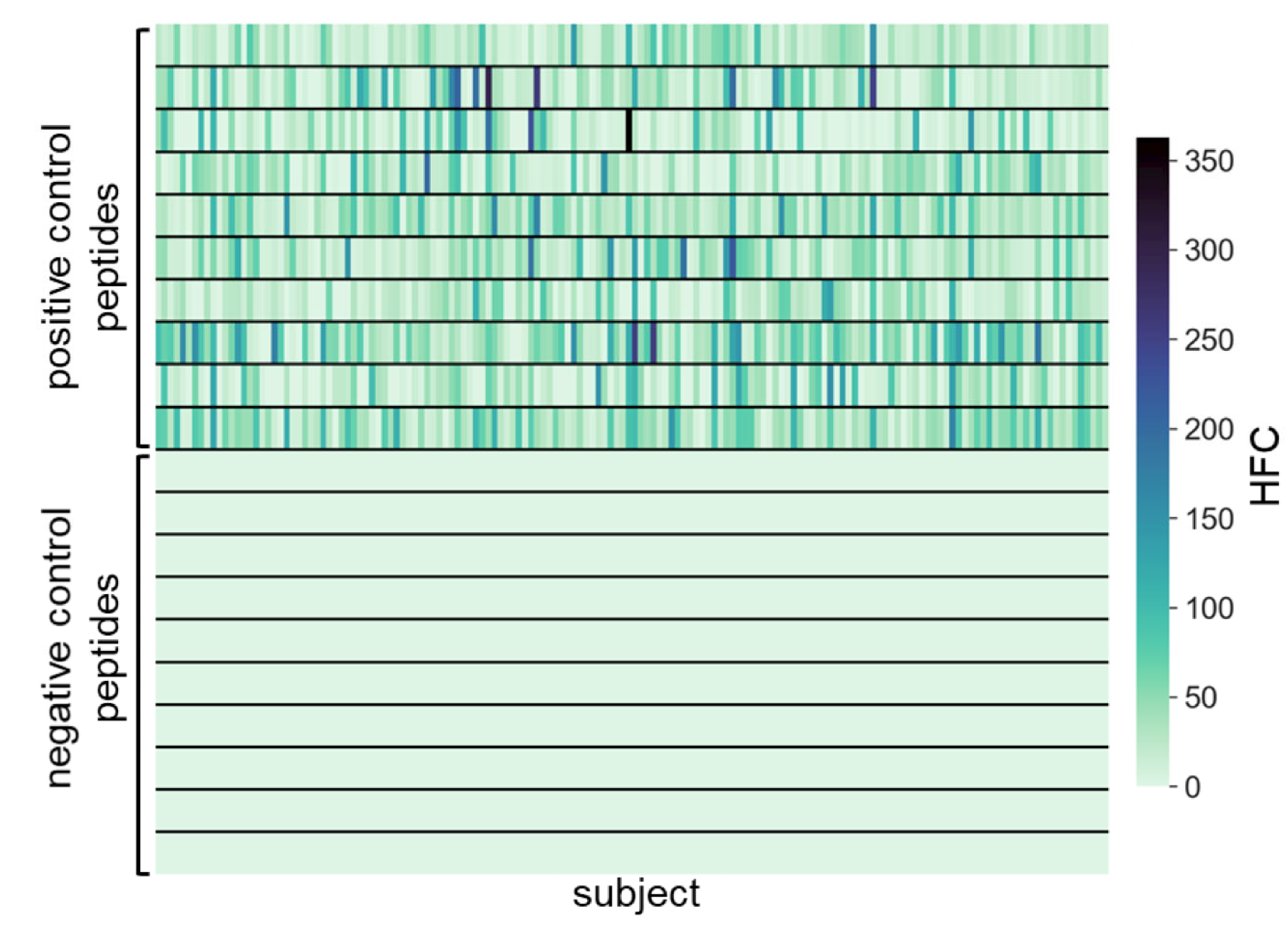
Positive and negative control peptide reactivities. Hits fold change values of positive and negative control peptides in serum samples from 156 subjects.

**Figure S3.**
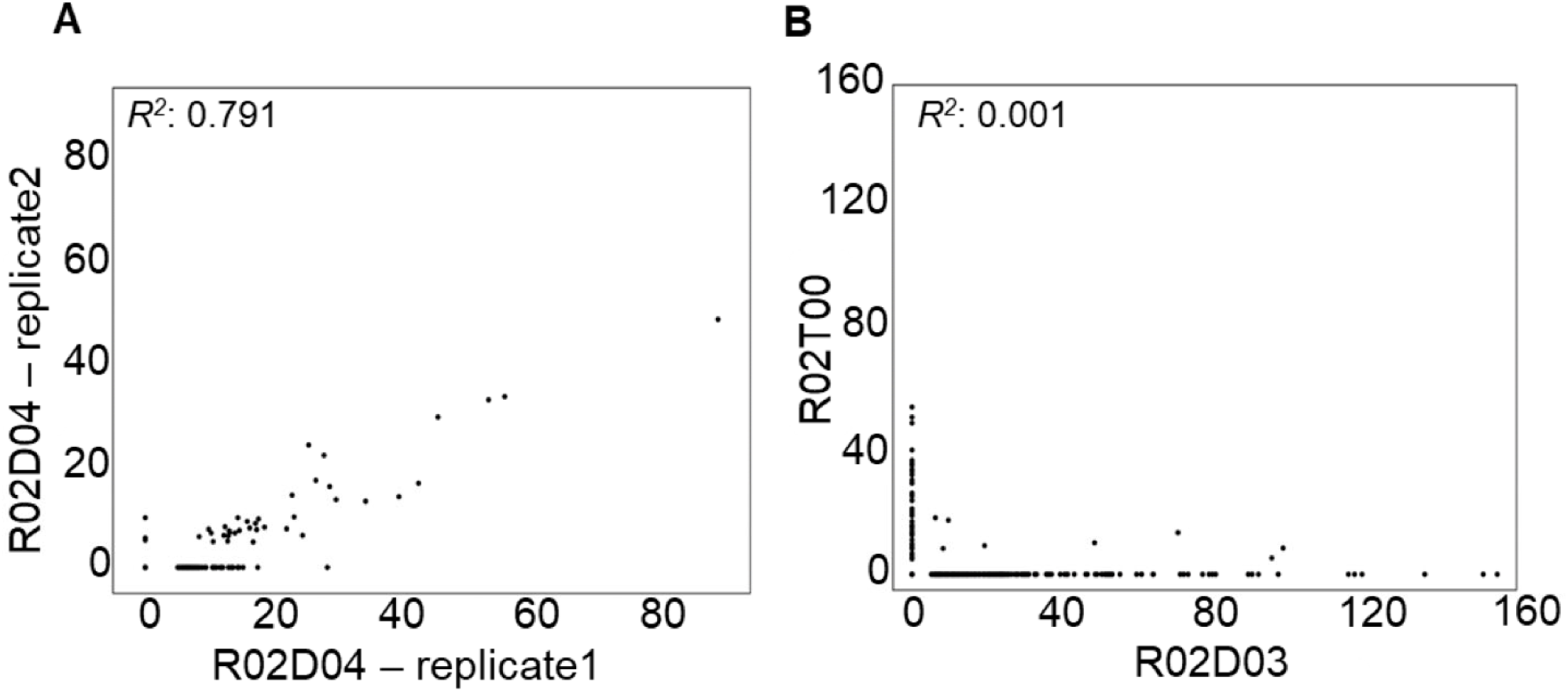
Reproducibility of the AnelloScan assay. Hits fold change values of AnelloScan peptides in **(A)** replicate and (B) non-replicate samples. “R02D03” and “R02D04” – serum samples of blood transfusion donors to recipient “R02”. “R02T00” – serum sample of recipient R02 before the transfusion event.

**Figure S4.**
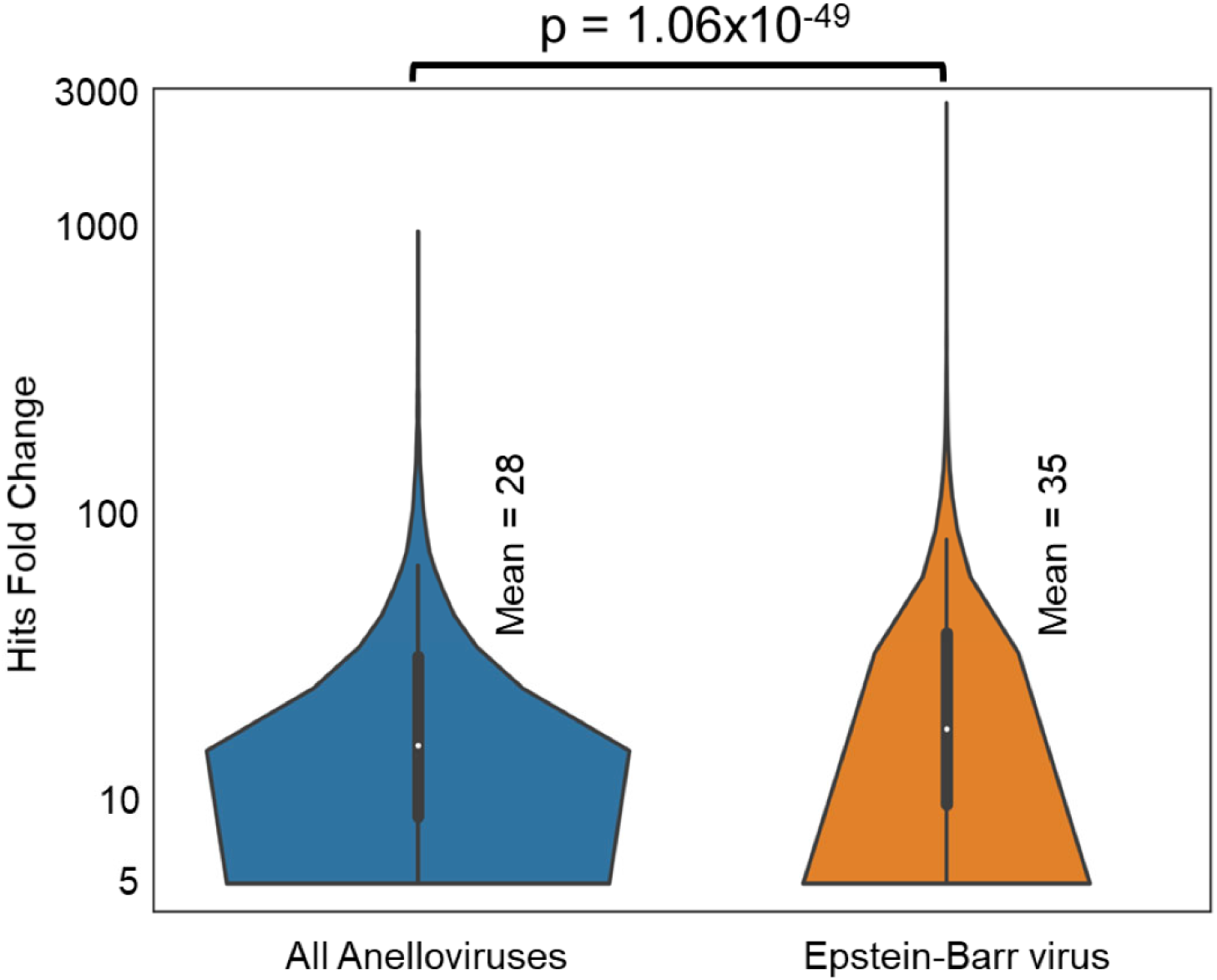
Anti-Anellovirus peptide versus Epstein-Barr virus peptide response strength. A violin plot showing the distribution of hits fold change values of all reactive AnelloScan peptides and EBV peptides from all individuals of the cross-sectional cohort (n=156).

**Figure S5.**
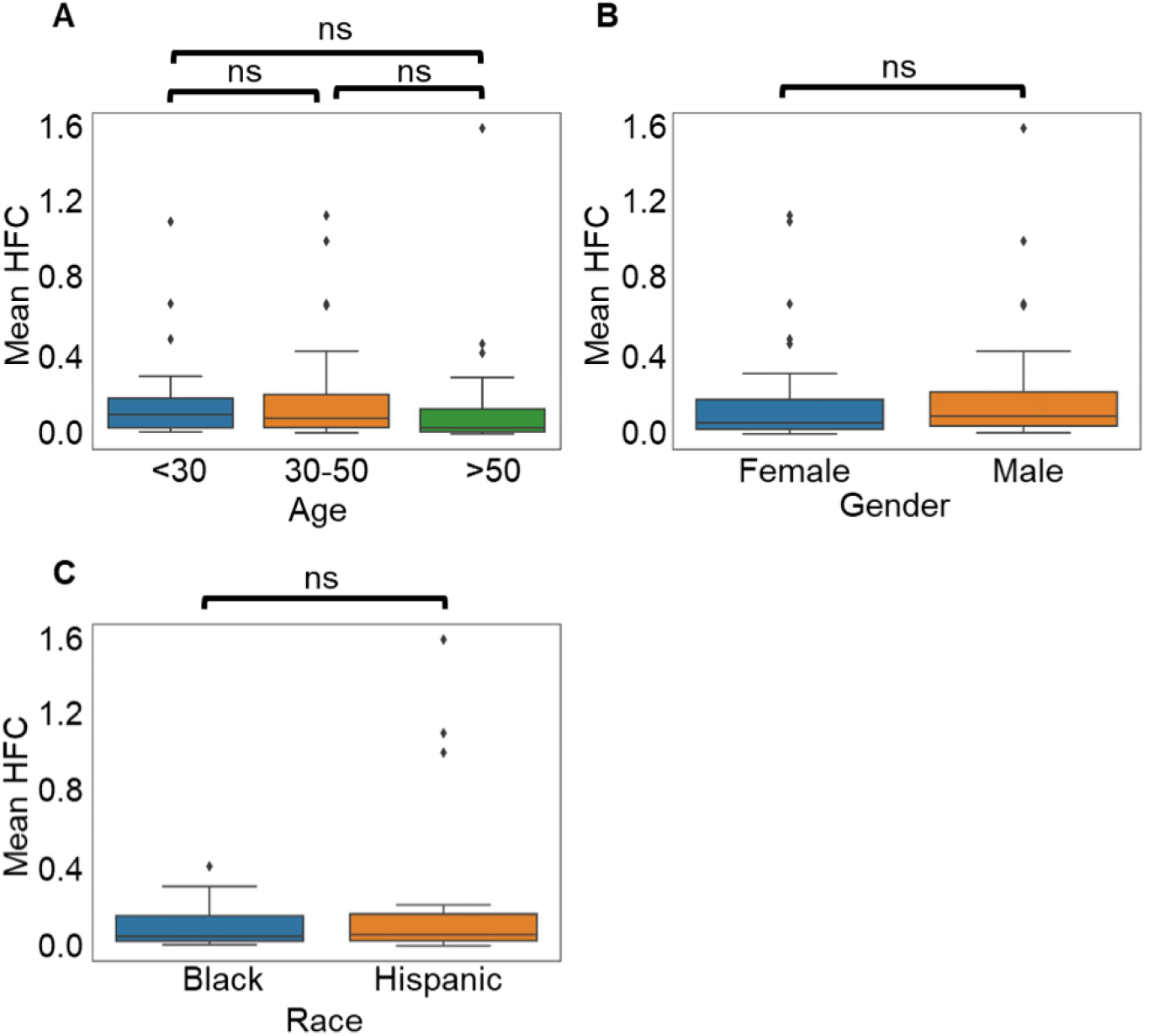
Demographic analysis of anellovirus antibodies. Mean antibody reactivities (Mean HFC) by **(A)** age, **(B)** gender and **(C)** race of the subject.

**Figure S6.**
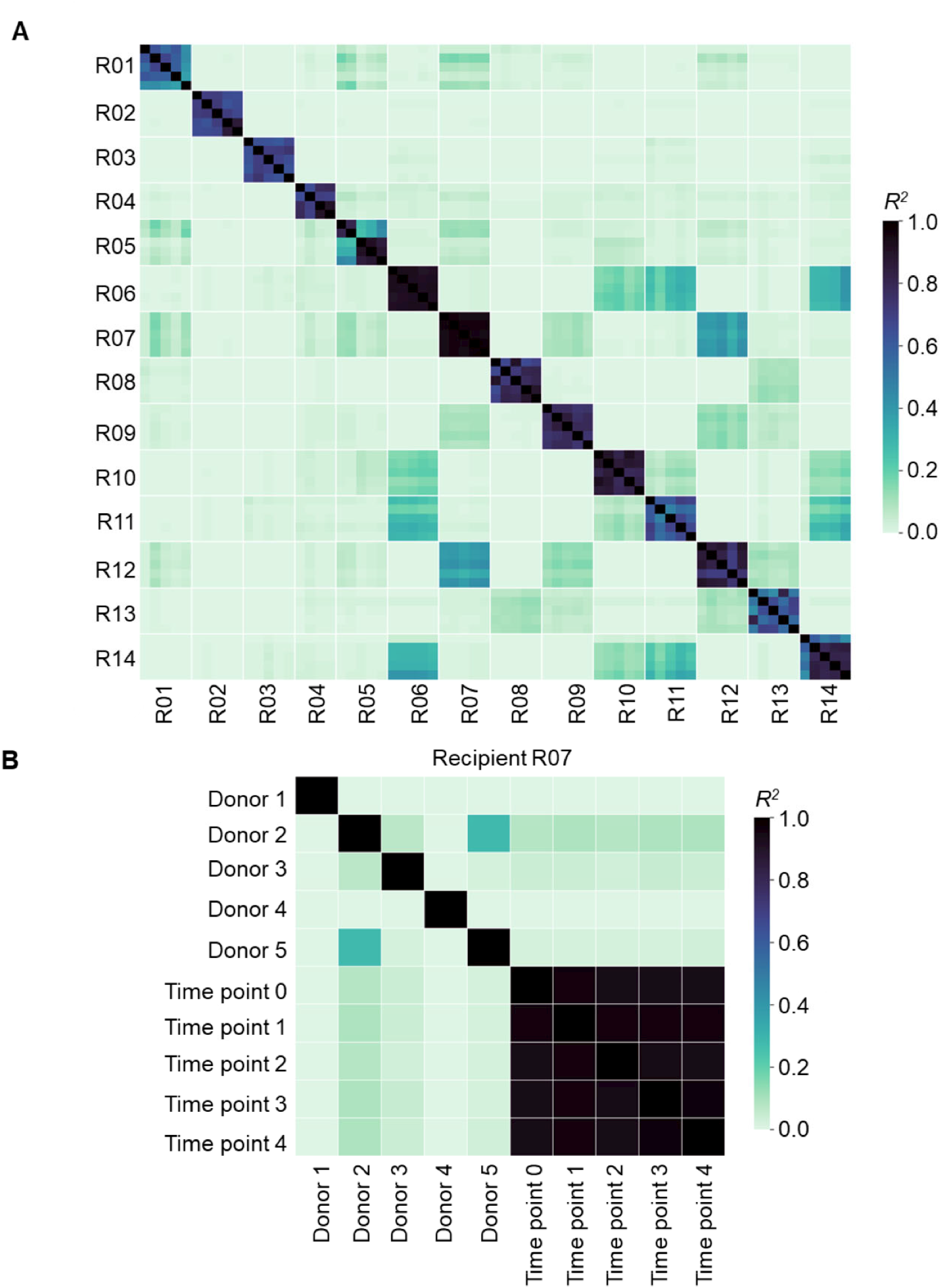
AnelloScan correlations are stable over time and not detectable in transfusion recipients. **A.** Anti-anellovirus antibody reactivities in a longitudinal cohort of 14 blood transfusion recipients at five time points. The heatmaps show the strength of the correlation of AnelloScan HFC values between two subjects. B. A correlation plot as in A shown for recipient R07 along with their 5 donors.

